# Effects of Prenatal Stress on Structural Brain Development and Aging in Humans

**DOI:** 10.1101/148916

**Authors:** Katja Franke, Bea van den Bergh, Susanne R. de Rooij, Tessa J. Roseboom, Peter W. Nathanielsz, Otto W. Witte, Matthias Schwab

**Author notes:** Corresponding author:* Dr. Katja Franke, Structural Brain Mapping Group, Department of Neurology, University Hospital Jena, Am Klinikum 1, D-07747 Jena, Germany.

## Abstract

Healthy brain aging is a major determinant of quality of life, allowing integration into society at all ages. Human epidemiological and animal studies indicate that in addition to lifestyle and genetic factors, environmental influences in prenatal life have a major impact on brain aging and age-associated brain disorders. The aim of this review is to summarize the existing literature on the consequences of maternal anxiety, stress, and malnutrition for structural brain aging and predisposition for age-associated brain diseases, focusing on studies with human samples. In conclusion, the results underscore the importance of a healthy mother-child relationship, starting in pregnancy, and the need for early interventions if this relationship is compromised.

## 1. Introduction

Human neurodevelopment begins *in utero* and continues through adolescence and early adulthood (Ernst and Korelitz, 2009). In the ‘Barker hypothesis’ / ‘Developmental Origins of Health and Disease (DOHaD) paradigm’ it has been suggested, that sensitive periods during human prenatal development are critical for the fetal origins of certain adult diseases, i.e. the fetus’ physiological adaptation to the (suboptimal) intrauterine environment has implications for health problems in later life which cannot completely be compensated for by later postnatal therapies (Barker, 1998). Plasticity and, hence, vulnerability of physiological systems is high during maturation, i.e. during fetal life. Thus, environmental influences are permanently modifying the function of physiological systems in critical phases of fetal development. More specifically, it has been proposed that prenatal environment, in particular, prenatal stress and malnutrition during gestation, trigger long-lasting modifications on the epigenome of the differentiating cell, thus resulting in changes in organ structure and adaptation of its metabolism, allowing optimal adaptation of the organism to its environment in order to ensure the immediate survival of the fetus (Barnes and Ozanne, 2011; Cao-Lei et al., this issue; Lillycrop and Burdge, 2011; Tarrade et al., 2015). However, the accumulation of oxidative stress due to adverse *in utero* conditions is suggested to consequently lead to accelerated cellular aging over the life course and therewith negatively affecting long-term (health) outcomes (Tarry-Adkins and Ozanne, 2014).

The brain is specifically susceptible to prenatal stress and malnutrition due to its long development and intrinsic plasticity, which is a basic property of the brain allowing learning and adaptation in later life (Bale, 2015; Griffiths and Hunter, 2014; Paus, 2005). In particular, the central nervous system (CNS) is gradually developing, including a complex timing of neuro- and gliagenesis, cell differentiation and migration, synaptogenesis. These highly coordinated processes have a high energy demand during development, with over half of the energy available is consumed by the brain during growth (Gibbons, 1998). The trajectory of neurodevelopment is critical for offspring quality of life (Rando and Simmons, 2015; Schuurmans and Kurrasch, 2013; Seckl and Holmes, 2007; Tarry-Adkins and Ozanne, 2014), with prenatal stress being widely linked to increased vulnerability for various cognitive, behavioral, and psychosocial problems (Charil et al., 2010; Talge et al., 2007; van den Bergh et al., this issue), and malnutrition during prenatal development compromising structural fetal cerebral development (Antonow-Schlorke et al., 2011) as well as being associated with developmental delays during childhood, impairments in life-long learning, cognitive deficits, behavioral and psychiatric disorders, as well as later-life neurodegenerative disorders (Brown et al., 2000; de Rooij et al., 2010; Faa et al., 2014; Hoeijmakers et al., 2014). After several prospective studies in human samples provided evidence for a strong link between antenatal stress and emotional and cognitive problems in the child, including increased risks of attentional deficit/hyperactivity disorder (ADHD), anxiety, and language delay (for a review please refer to Talge et al., 2007), it has been speculated that these links will be mediated by alterations in brain structure and function due to prenatal stress during fetal neurodevelopment.

A wealth of animal studies demonstrated that prenatal stress is indeed affecting the morphology of the offspring’s brain, usually showing reduced tissue volumes on both, macroscopic and microscopic levels (for a review please refer to Charil et al., 2010). Experimental animal studies also provided strong evidence for malnutrition during prenatal life causing widespread disturbances of early organizational processes in cerebral development that result in permanent impairments in brain structure and function (Antonow-Schlorke et al., 2011; Benton and a.i.s.b.l, 2008; Grantham-McGregor and Baker-Henningham, 2005; Keenan et al., 2013; Morgane et al., 1993; Morley and Lucas, 1997; Muller et al., 2014; Olness, 2003; Rodriguez et al., 2012; Wainwright and Colombo, 2006; Walker et al., 2007), subsequently determining altered postnatal cognitive and behavioral performances (Keenan et al., 2013; Rodriguez et al., 2012). However, the effects of environmental influences animal studies are not easily transferable to the human situation. Most developmental programming studies have been mainly conducted in polytocous, altricial rodents, which show substantially different trajectories of fetal and neonatal brain development from monotocous, precocial mammals, including humans (Fontana and Partridge, 2015; Ganu et al., 2012). Thus, more studies on prenatal stress and brain development in human samples were consequentially called for (Charil et al., 2010).

Therefore, the scope of this review is to summarize all studies examining the effects of a suboptimal fetal environment on brain structure in human samples, covering age ranges from infants up to late adulthood. We discuss studies based on anatomical and neuroimaging data, revealing neuroanatomical differences in offspring’ brain associated with maternal stress, maternal anxiety, or maternal depression during pregnancy as well as with fetal malnutrition. We try to dissect the extent of neurostructual abnormalities in relation to the prenatal environmental influence, the age at which they become apparent, and the relation of neurostructural abnormalities to cognitive, neuropsychiatric and mental disorders.

## 2. Studies examining neuroanatomical differences associated with maternal stress, anxiety, and depression during pregnancy

The first studies examining the effects of maternal stress, anxiety, or depression during pregnancy on brain development have used anatomical measures, e.g. head circumference at birth. Studies analyzing neuroimaging data were using diverse magnetic resonance imaging (MRI) modalities, pulse sequences, and protocols as well as a variety of analytical techniques for processing MRI data in order to cover and quantify the multidimensional aspects of changes in brain structure throughout the lifespan. These data may result in several different markers for the individual brain structure that are sensitive to and capture shared as well as marker-specific information on brain tissue changes (Cherubini et al., 2016; Groves et al., 2012). The most widely used modality is T1-weighted MRI, providing a number of parameters, including voxel-wise, regional and global gray matter (GM) and white matter (WM) volumes, volumetric data for subcortical regions, cortical thickness, and cortical surface area. Complementary measures of individual brain characteristics provide information about the myelin content and white matter integrity in regions of interest (ROIs) and fiber tracts (from T2-weighted MRI), and information about fiber density, axonal diameter, myelination in white matter, as well as several measures of water diffusion in brain tissue (from diffusion tensor imaging; DTI). For a short overview of the different modalities and resulting parameters please refer to Table 1 in Franke et al. (this issue).

**Table 1.**
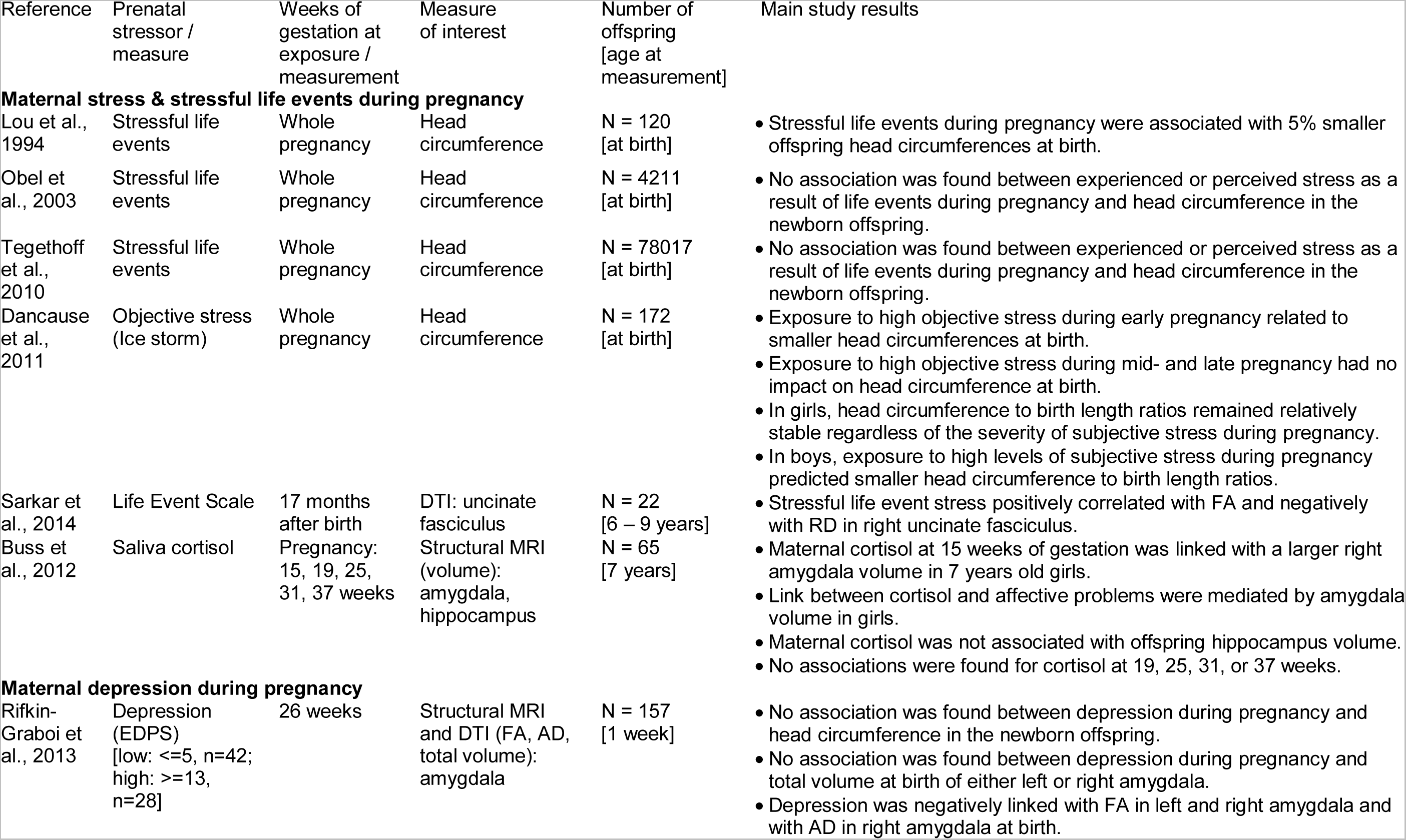
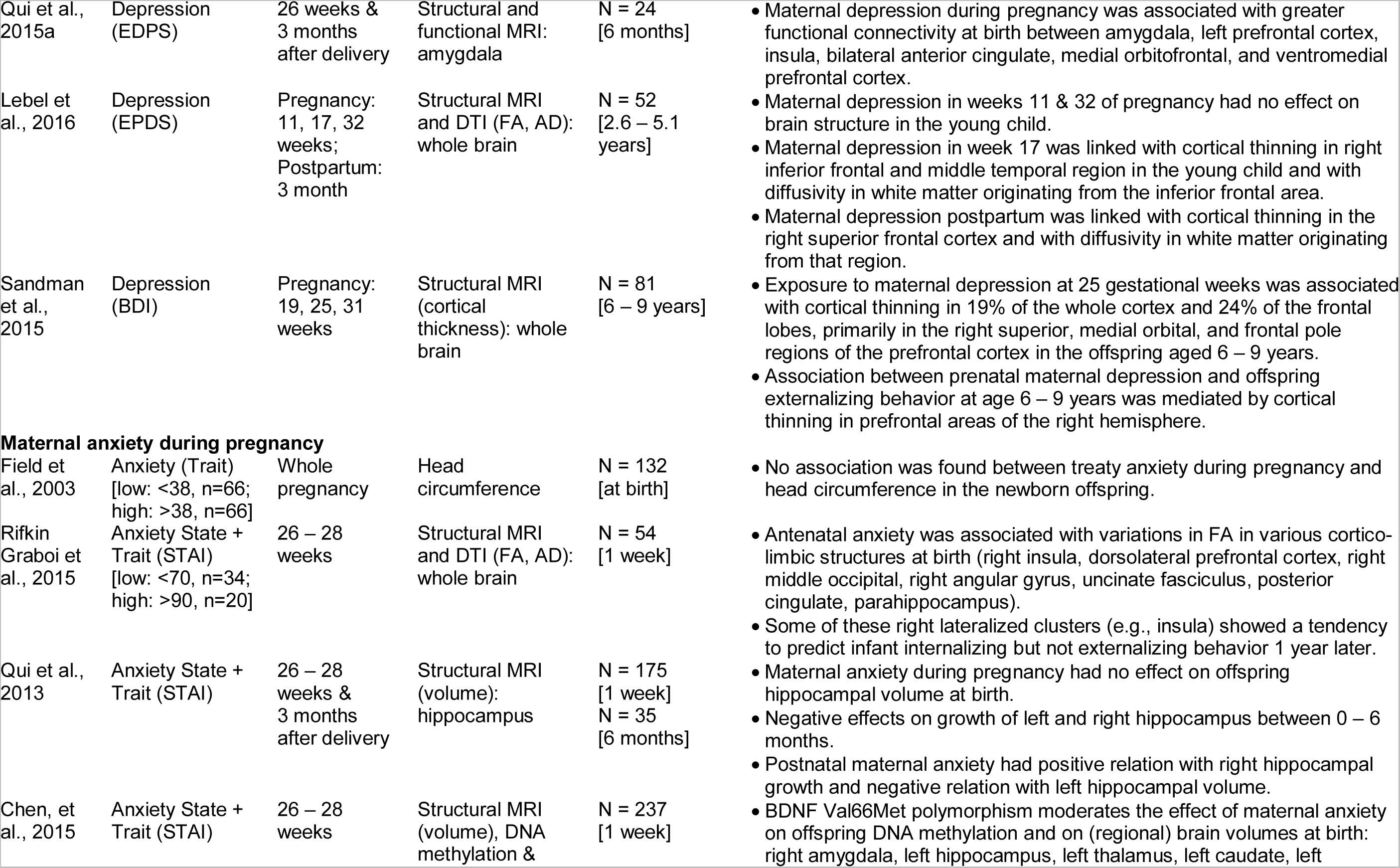
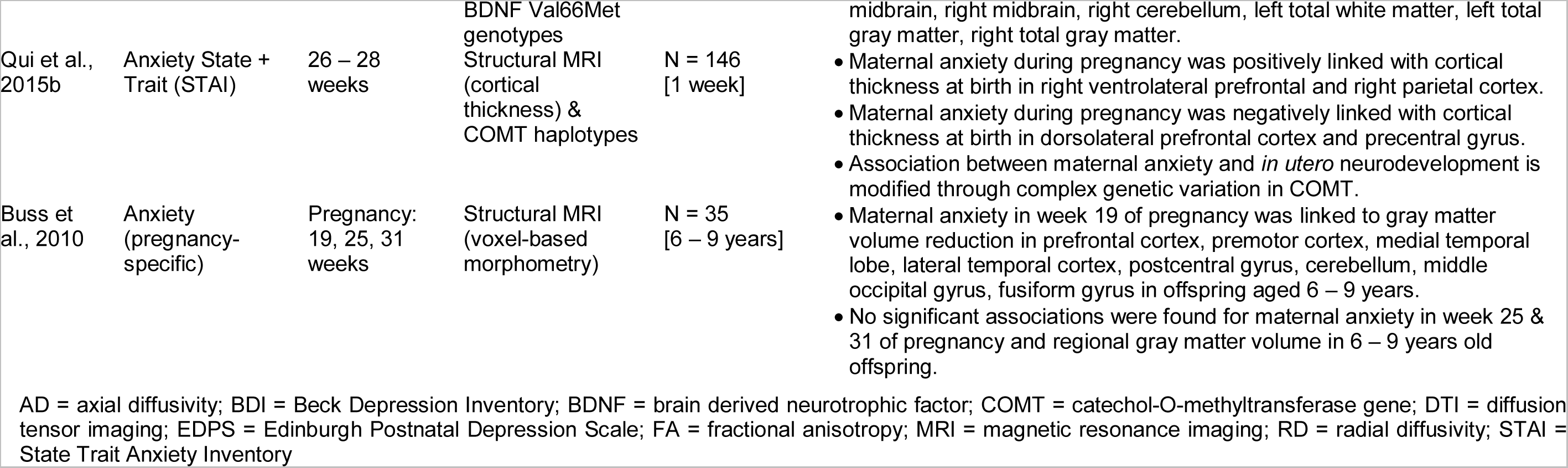
Maternal stress, anxiety, depression during pregnancy and offspring neuroanatomical brain correlates

A condensed overview of all studies exploring the associations between maternal stress, anxiety, or depression during pregnancy and neuroanatomical measures in neonates, infants, preschool-, and school-aged children con be found in Table 1.

### 2.1 Effects of maternal stress during pregnancy

Using anatomical birth measures, a population-based study from Denmark reports that stressful life events (e.g., separation, job loss, theft, death of a spouse or partner) during pregnancy is negatively associated with offspring head circumference at birth, corresponding to a reduction of at least a 5% in brain volume at birth among offspring of mothers who experienced stressful life-events during pregnancy. The authors concluded that these findings suggest the existence of a fetal stress syndrome, which adversely effects fetal brain development (Lou et al., 1994).

However, two other population-based studies from Denmark (Obel et al., 2003; Tegethoff et al., 2010) could not find any association between stressful life events during pregnancy and offspring head circumference at birth, although Obel et al. (2003) reported similar values of head circumferences in the newborn offspring of mothers who experienced stressful life events during pregnancy as were reported in the study by Lou et al. (1994).

Another study using anatomical birth measures explored the effects of timing and severity of exposure during pregnancy to objective and subjective stress due to a natural disaster (i.e. Ice storm) on fetal development. The results showed smaller head circumferences at birth being associated with high levels of objective stress levels during early pregnancy but not during in mid- and late pregnancy, while controlling for other variables such as age, obstetric complications, socioeconomic status, and trait anxiety. Additionally, exposure to high levels of subjective stress during pregnancy predicted smaller head circumference to birth length ratios in boys, whereas in girls, head circumference to birth length ratios remained relatively stable regardless of the severity of subjective stress during pregnancy (Dancause et al., 2011).

Sarkar et al. (2014) reported results of a small sample of women (*n* = 22) that underwent amnioscentesis and completed a life event questionnaire on stressful life events during pregnancy retrospectively, at 17 months after birth. Maternal self-report of stress during pregnancy was associated with differences in the development of white matter within brain networks underlying child social behavior in the 6 – 9 years old offspring. These changes in WM microstructure were suggested to reflect hypermyelination in the right uncinate fasciculus, which links the limbic region with the prefrontal cortex (Sarkar et al., 2014).

Another prospective study utilized cortisol as a physiologically biomarker for stress to examine the association of maternal cortisol in early, mid-, and late gestation with subsequent measures of amygdala and hippocampus volume and affective problems in the 7 years old offspring (Buss et al., 2012). Higher maternal cortisol levels in earlier but not later gestation were found to be associated with a larger right amygdala volume in girls, but not in boys. Furthermore, higher maternal cortisol levels in early gestation were associated with more affective problems in girls, being in part mediated by amygdala volume. Thus, maternal stress hormone levels in pregnancy was linked with subsequent child amygdala volume and affect. The results underscore the importance of the intrauterine environment and suggest the origins of neuropsychiatric disorders may have their foundations early in life (Buss et al., 2012).

### 2.2 Effects of maternal depression during pregnancy

One of the most important studies with originally 1176 subjects is the prospective longitudinal study “Growing up in Singapore towards Healthy Outcomes” (GUSTO; Soh et al., 2014). Pregnant mothers were recruited at 13 weeks of pregnancy. Maternal depression and anxiety were measured at 26 – 28 weeks of pregnancy and 3 months after pregnancy. A total of 388 infants underwent brain MRI scans one week after birth. Out of them, 30 infants got a 2^nd^ MRI scan at 6 weeks of age and another 50 infants got a 2^nd^ MRI scan at 6 months of age.

Using data from the GUSTO project, Rifkin-Graboi et al. (2013) neither observed differences in head circumference at birth, nor for volume of the amygdala 1 week after birth between offspring of mothers with high vs. low depression in pregnancy. However, the study reported a negative association between maternal depression in pregnancy and white matter fiber measures in the right amygdala, a brain region closely associated with stress reactivity and vulnerability for mood anxiety disorders. According to the authors these findings suggest the prenatal transmission of vulnerability for depression from mother to child (Rifkin-Graboi et al., 2013).

At 6 months of age, maternal depression was related to greater functional connectivity between brain regions that are involved in the activation and regulation of emotional states, i.e., amygdala, left prefrontal cortex, insula, bilateral anterior cingulate, medial orbitofrontal and ventromedial prefrontal cortices in the offspring, which are largely consistent with connectivity patterns observed in adolescents and adults with major depressive disorder. Thus, neuroimaging correlates of the prenatal transmission of phenotypes associated with maternal mood are already apparent in infants at 6 months of age (Qiu et al., 2015a).

A recent Canadian study reported that women’s depression in the 2^nd^ trimester of pregnancy was associated with cortical thinning in right inferior frontal and middle temporal region as well as with white matter measures of fibers emanating from the inferior frontal area in preschool children (Lebel et al., 2016). However, the associations between maternal depression in gestational week 17 and white matter fiber measures were no longer significant after correction for postpartum depression. Additionally, postpartum depression was also related with cortical thinning with children’s right superior frontal cortical thickness and with white matter measures of fibers originating from that region; the effect remained significant after correction for prenatal maternal depression. Maternal depression in gestational weeks 11 & 32 was not significantly related with altered gray or white matter structures (Lebel et al., 2016).

Another prospective study examined the association between fetal exposure to maternal depressive symptoms and cortical thickness in the 6 – 9 years old offspring (Sandman et al., 2015). Maternal reports of depressive symptoms were associated with cortical thinning, showing the strongest association with maternal depression at 25 weeks’ gestation. The largest associations were in the prefrontal, medial postcentral, lateral ventral precentral and postcentral regions of the right hemisphere. Moreover, cortical thinning in prefrontal areas of the right hemisphere mediated the association between maternal depression and child externalizing behavior. Remarkably, the observed pattern of cortical thinning seems to be similar to patterns in children, adolescents and depressed patients. The authors suggest the observed cortical thinning in children born to mothers with higher depressive symptoms during pregnancy probably reflecting accelerated brain maturation (Sandman et al., 2015).

### 2.3 Effects of maternal anxiety during pregnancy

Using anatomical birth measures, trait anxiety in the mothers during pregnancy was not found to have an effect on the neonate’s head circumference (Field et al., 2003).

Using data from the GUSTO project, and examining the effects of maternal anxiety during pregnancy on the brain structure in the newborn offspring resulted in variations in white matter fiber measures in dorsolateral prefrontal cortex and right insula (regions important to cognitive-emotional responses to stress), right middle occipital (sensory processing) and the right angular gyrus, uncinate fasciculus, posterior cingulate parahippocampus (social cognition, social-emotional functioning). Additionally, the authors reported a tendency for some of these lateralized clusters predicting infant internalizing but not externalizing behavior at 1 year of age (Rifkin-Graboi et al., 2015).

Prenatal maternal anxiety was also found to have no effect on bilateral hippocampal volume at birth. However, maternal anxiety was negatively correlated with growth of both the left and right hippocampus over the first 6 months of life. Furthermore, a positive association was observed between postnatal maternal anxiety and offspring right hippocampal growth and a negative one between postnatal maternal anxiety and left hippocampal volume at 6 months of age (Qiu et al., 2013).

The brain derived neurotrophic factor (BDNF) Val66 gene is known to underlie the synaptic plasticity throughout the central nervous system (Kleim et al., 2006; McHughen et al., 2010; Wang et al., 2014). Furthermore, the BDNF Val66Met polymorphism was found to moderate the impact of environmental conditions on brain-based phenotypes (Gunnar et al., 2012; Hayden et al., 2010; Juhasz et al., 2011; Mata et al., 2010; Willoughby et al., 2013). Chen et al. (2015) examined the association between neonatal DNA methylation and brain substructure volume, as a function of BDNF genotype. The results showed that infant, but not maternal, BDNF genotype strongly influences the association between antenatal anxiety and the offspring epigenome at birth as well as that between the epigenome and neonatal brain structure, especially with amygdala and hippocampus volumes. More specifically, in the Met/Met polymorphism group there was a greater impact of antenatal maternal anxiety on the infant DNA methylation then in both other groups. The authors concluded that these findings suggest that differential susceptibility to specific environmental conditions may be both tissue and function specific (Chen et al., 2015).

Qiu et al. (2015b) observed an effect of maternal anxiety during pregnancy on neonatal frontal cortical thickness that was moderated by functional variants of the catechol-O-methyltransferase (COMT) gene, which regulates catecholamine signaling in the prefrontal cortex and is implicated in anxiety, pain, and stress responsivity. The A-val-G (AGG) haplotype moderated the positive link between maternal anxiety and thickness of the right ventrolateral prefrontal, right parietal cortex and precuneus. The G-met-A (GAA) haplotype modulated the negative link between maternal anxiety and thickness of the dorsolateral prefrontal cortex and bilateral precentral gyrus.

In a prospective study examining the effect of the temporal pattern of pregnancy anxiety to specific changes in brain morphology, Buss et al. (2010) reported an association between pregnancy-specific anxiety at week 19 of gestation and gray matter volume reductions extending from prefrontal to occipital regions in the 6 – 9 years old offspring. However, high pregnancy anxiety at 25 and 31 weeks gestation was not significantly associated with local reductions in gray matter volume. The authors thus suggested that altered gray matter volume in brain regions affected by prenatal maternal anxiety may render the developing individual more vulnerable to neurodevelopmental and psychiatric disorders as well as cognitive and intellectual impairment (Buss et al., 2010).

## 3. Neuroimaging studies revealing neuroanatomical differences associated with malnutrition during pregnancy

A severe form of prenatal stress is constituted by prenatal malnutrition. The developing brain is highly dependent on the availability of nutrients and a lack of sufficient nutrition forms a serious threat to normal brain development (Ramel and Georgieff, 2014). There are few studies in humans in which the direct effects of prenatal malnutrition on neuroanatomy have been measured. Quite a number of studies have investigated the associations between small size at birth and brain morphology though. Size at birth is an indirect measure for the fetal environment and small size at birth can result from prenatal malnutrition due to maternal malnutrition, placental insufficiency, extreme maternal vomiting or a multiple pregnancy. First results in twin studies suggested greater birth weight to be associated with a sustained and generalized increase in brain volume and cortical surface area that persist into late adolescence. Furthermore, these surface area alterations were not only related to cognition, but also to the risk of several mental disorders (Raznahan et al., 2012). Very small size at birth (e.g. very low birth weight or extremely low birth weight) usually results from being born (very) preterm. Below we will give an overview of studies in which associations between birth weight and neuroanatomy were studied (excluding studies of very low birth weight, preterm birth or multiple pregnancies).

Sanz-Cortes et al. (2014) performed MR imaging in fetuses at 37 weeks of gestation and compared 54 small for gestational age (SGA, defined as birth weight below the 10th percentile adjusted for gestational age) babies to 47 babies with normal size for age (Sanz-Cortes et al., 2014). They found that the brain stem and cerebellum were smaller in the SGA group and this also correlated with neurobehavioral outcomes. A number of other studies investigated brain-imaging outcomes in adolescents and young adults who were born SGA. The group of Martinussen et al. conducted MRI studies in 15-year old boys and girls: A first study showed reduced brain volume in those born SGA compared to controls and a second study demonstrated smaller brains and proportionally smaller regional brain volumes, while hippocampus volume was a significant predictor of cognitive function (Martinussen et al., 2005; Martinussen et al., 2009). In a third study, they again showed smaller brain volumes in 15-year olds born SGA, but these findings were only present in those with fetal growth restriction as evidenced by frequent ultrasounds in pregnancy (Rogne et al., 2015). An additional study, performed at age 20 years, showed regional reductions in cortical surface area and total brain volume, cortical GM, cerebral WM and putamen volumes were reduced (Ostgard et al., 2014). The most marked reduction was seen in those with fetal growth restriction. These findings largely coincide with findings by de De Bie et al. (2011) who investigated 4-to 7-years old children born SGA and who displayed reduced cerebral and cerebellar grey and white matter volumes, smaller volumes of subcortical structures and reduced cortical surface area. Regional differences in prefrontal cortical thickness suggested a different development of the cerebral cortex.

Other studies compared groups of people born with low birth weight (LBW < 2000 grams) to people born with normal weights. Odberg et al. (2010) showed an increased risk for 19-year olds born with LBW to have prominent ventricles, global loss of white matter and thinning of the corpus callosum. A specific analysis of the corpus callosum, showed that specifically the posterior third subregion was smaller (Aukland et al., 2011). At age 15 – 16 years, LBW was found to be associated with a smaller surface area of the lateral and medial orbitofrontal cortex, and right inferior frontal gyrus. This study also showed that a lower caudate volume mediated the effect of low birth weight on impaired inhibitory control (Schlotz et al., 2014).

The importance of gestational age interacting with birth weight was demonstrated in a study of 157 6-years old boys from a normal birth weight spectrum, where these interactive effects predicted caudate volumes and shapes (Qiu et al., 2012). Boys with relatively low birth weight and shorter gestation had smaller caudate volumes, reflected by shape contraction in the middle body. They also performed worse in a motor response task. The authors concluded that prenatal influences on neurocognitive and brain development are not limited to the extreme range, but occur across the entire population.

Finally, a unique study was performed in a very large group of 1254 older adults with a mean age of 75 years (Muller et al., 2014). Findings indicated that lower ponderal index at birth (a measure for body weight/height at birth) was associated with smaller volumes of total brain volume and total white matter. Additionally, lower ponderal index was associated with slower processing speed and reduced executive functioning but interestingly this was only true in those with low education.

Overall, these studies provide evidence that small size at birth is associated with altered brain morphology during gestation, in childhood, adolescence and well into older age. Total brain volume is mostly found to be reduced, with (regional) volume differences also relating to clinical differences in cognitive function. As said though, size at birth is only an indirect reflection of the fetal environment. Evidence from the Dutch famine birth cohort study has importantly shown that an adverse prenatal environment, especially in the beginning of pregnancy, is associated with a range of adverse health effects that were not dependent on birth weight (Roseboom et al., 2006). Those exposed to the famine in early gestation were even found to have normal size at birth. The Dutch famine was a five-month period at the end of World War II during which the western part of the Netherlands was struck by a severe famine. The famine was of course a humanitarian disaster, but the fact that it was a relatively short period of severe malnutrition in a population well fed before and after the famine makes it a unique opportunity to study consequences of prenatal exposure to malnutrition. Direct effects of the famine on the development of the central nervous system (CNS) were shown by a study by Stein et al. (1975) who reported that babies who had been exposed to the Dutch famine during the first gestational trimester showed an increase in the prevalence of congenital anomalies of the CNS, including spina bifida and hydrocephalus. The effects of exposure to malnutrition in early gestation on brain development were not limited to short term effects though. Two different studies have demonstrated that these effects can still be observed in middle and also late life. The first study was a small study that was performed in a group of 18 schizophrenic patients who were matched to 18 healthy controls at the age of 51 years (Hulshoff Pol et al., 2000). MRI findings showed that intracranial volume was decreased in famine exposed schizophrenia patients and prenatal famine exposure alone was related to an increase in brain abnormalities, predominantly white matter hyperintensities. A second study was performed in a subsample of 118 members of the Dutch famine birth cohort, which is a cohort of people who were all born in a local hospital in Amsterdam around the time of the famine (de Rooij et al., 2016). At the age of 68 years, MR scans of a group of 41 men and women exposed to the famine in early gestation were compared to those of a group of 77 people unexposed to the famine *in utero*. The results showed that prenatally famine exposed males but not females, had a smaller intracranial and total brain volume compared to unexposed subjects. The smaller brain volume found in exposed men may have been the consequence of an early interruption of brain development caused by nutritional deficiency in the first trimester of gestation, which is still visible after 68 years. In line with this explanation, a study in 256 children at the age of 6 – 8 years showed that a specific nutritional deficiency, maternal folate insufficiency in early pregnancy, was also associated with a reduction in total brain volume, which was also related to poorer cognitive performance (Ars et al., 2016). Alternatively, the brains of famine-exposed men may have been more vulnerable to the effects of ageing. Additional analyses, utilizing the innovative *BrainAGE* approach, which was designed to indicate deviations in age-related spatiotemporal brain changes, have demonstrated that prenatally famine exposed elderly males also show advanced brain aging by about 4 years (Franke et al., 2017). A third option would be that both early structural and late ageing effects have played a role in the smaller brain volumes after prenatal famine exposure. The folate and famine studies differed with regards to the sex differences found in the famine study, which were not shown in the folate study. An explanation for this may be that the Dutch famine birth cohort subsample study was hampered by selective participation of more healthy females, as it was previously shown that women exposed to the famine in early gestation had increased mortality up to the age of 63 years (van Abeelen et al., 2012). However, it could also be that the development of the CNS in men is more vulnerable to effects of malnutrition. Ivanovic et al. (2000) demonstrated in a study in Chilean children that malnutrition in the first year of life had a clear effect on brain volume at age 18 in both males and females, but the effect in males was much larger than the effect in females.

In summary, the prenatal malnutrition studies are generally in line with the results from studies that investigated small size at birth in relation to brain morphology. Prenatal malnutrition seems to have a clear global effect on brain structure in that it is associated with overall brain volume and not so much with differences in specific regions. What the exact clinical implications of these findings are, especially in older age, remains to be elucidated.

## 4. Conclusion

Several significant associations were found between maternal stress, anxiety and depression in the pregnant mother and changes in the offspring brain structure brain, inducing larger amygdala volume in 7 years old girls (Rifkin-Graboi et al., 2013), smaller hippocampal growth in infants (Qiu et al., 2013), GM volume reduction in several cortical areas at birth (Buss et al., 2010; Chen et al., 2015), changes in cortical thickness of prefrontal areas at birth (Qiu et al., 2015b), in young childhood (Lebel et al., 2016), as well as at age 6 – 9 years (Sandman et al., 2015), structural changes in several prefrontal cortical and cortico-limbic structures in infants (Rifkin-Graboi et al., 2015), and aberrations in subcortical WM structures in infants (Rifkin-Graboi et al., 2013). These structural changes found are assumed to potentially mediate the link between maternal stress in pregnancy and offspring cognitive, behavioral and emotional problems, thus as markers of prenatal stress, which are increasing the risk for mental health problems. Empirical evidence for these effects being moderated by genetic differences and mediated by epigenetic changes begins to being revealed in human studies (Chen et al., 2015; Rifkin-Graboi et al., 2015). These results underscore the importance of a healthy mother-child relationship starting in pregnancy and the need for early interventions if this relationship is compromised.

Previously, prenatal stress and fetal malnutrition was widely linked to increased vulnerability for various cognitive, behavioral, and psychosocial problems, as well as later-life neurodegenerative disorders (Brown et al., 2000; Charil et al., 2010; de Rooij et al., 2010; Faa et al., 2014; Hoeijmakers et al., 2014; Talge et al., 2007). Experimental animal studies convincingly demonstrated that prenatal stress and fetal malnutrition is causing widespread disturbances of early organizational processes in the offspring’s cerebral development that result in permanent impairments in brain structure and function affecting the morphology on both, macroscopic and microscopic levels (Antonow-Schlorke et al., 2011; Benton and a.i.s.b.l, 2008; Charil et al., 2010; Grantham-McGregor and Baker-Henningham, 2005; Keenan et al., 2013; Morgane et al., 1993; Morley and Lucas, 1997; Muller et al., 2014; Olness, 2003; Rodriguez et al., 2012; Wainwright and Colombo, 2006; Walker et al., 2007). This review aimed to wrap up the latest research in humans on the effects of maternal stress and fetal malnutrition during gestation on neurodevelopment and lifelong consequences on brain structure in the offspring.

Most studies utilizing non-invasive imaging markers for brain structure have found associations between maternal stress, cortisol, anxiety, or depression during pregnancy and structural brain changes in the offspring, including smaller hippocampal growth effects, larger amygdala volume, GM volume reduction in several cortical areas, cortical thinning in several prefrontal areas, and microstructural changes in prefrontal cortical and cortico-limbic structures as well as subcortical areas, hypermyelination in certain WM structures, in part probably reflecting accelerated brain maturation (Sandman et al., 2015). Consequently, these structural changes are assumed to mediate the link between maternal distress in pregnancy and offspring cognitive, behavioral and emotional problems, finally increasing the risk for mental health problems, being mediated by gene x environment interactions and gene x epigenome interactions (Chen et al., 2015; Rifkin-Graboi et al., 2015). More specifically, alterations relating to maternal anxiety were found in the offspring’s brain structure determining cognitive-emotional responses to stress, sensory processing, and social-emotional functioning (Rifkin-Graboi et al., 2015). Moreover, these patterns of altered brain structure observed in offspring experiencing maternal stress during gestation resemble of those observed in depressed patients (Sandman et al., 2015). Additionally and in line with the assumption of gender-specific mechanisms of maturation, aging, disease affection etc. (Aiken and Ozanne, 2013; Azad et al., 2007; Grossi et al., 2005; Pinn, 2003), sex-specific effects of prenatal stress on brain maturation and aging started to reveal (Buss et al., 2010). However, in the only paper reporting correction for multiple testing, the associations between depression in pregnancy and alterations of neonate brain regions vanished (Rifkin-Graboi et al., 2015).

Lack of sufficient nutrition during gestation constitutes a severe form of prenatal stress, being related to increased health problems and disturbances in brain development (Lillycrop and Burdge, 2011; Ozanne and Hales, 2004; Ramel and Georgieff, 2014; Tarry-Adkins and Ozanne, 2014). Only few studies in humans have directly measured the effects of prenatal malnutrition on neuroanatomy, instead investigating the associations between brain morphology and size at birth, which is an indirect measure for the fetal environment, with small size at birth resulting from prenatal malnutrition due to maternal malnutrition, placental insufficiency, extreme maternal vomiting or a multiple pregnancy. Studies examining intrauterine growth restriction and small size at birth are repeatedly reporting altered brain morphology during gestation, in childhood, adolescence and well into older age. These alterations, including smaller total and regional brain volumes, reductions in cortical surface area and prefrontal cortical thickness, have been demonstrated to correlate with neurobehavioral outcomes and impaired cognitive function, like slower processing speed and reduced executive functioning. However, evidence from the Dutch famine birth cohort study has shown that an adverse prenatal environment, especially in the 1^st^ trimester of pregnancy, is associated with an increase in brain abnormalities that were not dependent on birth weight, including smaller intracranial and total brain volume as well as advanced brain aging in late adulthood, suggesting early interruption of brain development caused by nutritional deficiency or increased vulnerability to the effects of aging, or both. Being in line with developmental programming models suggesting that long-term (health) outcomes of adverse *in utero* conditions will be more prominent in male than female offspring (Aiken and Ozanne, 2013), these effects were only shown in men. In summary, studies of prenatal malnutrition in humans show global effects on brain structure. However, the exact clinical implications of these findings, especially in older age, remain to be elucidated.

In conclusion, maternal stress during gestation and prenatal malnutrition are associated with gender-specific disturbances in early neurodevelopment, resulting in lifelong alterations of brain morphology in the offspring and consequently cognitive, behavioral and emotional problems, finally increasing the risk for mental health problems. Future work should further explore how epigenetic and environmental interactions are mediating the lifelong effects of several factors of maternal stress during pregnancy (e.g. malnutrition, maternal obesity and diabetes, smoking during pregnancy, twin pregnancy, placental insufficiency, anxiety) on neuroanatomical maturation and aging in order to identify subtle, yet clinically-significant, changes in brain structure, thus contributing to a better understanding of the consequences of prenatal environment on life-long brain health as well as to an early diagnosis of neurodegenerative diseases and facilitating early treatment or preventative interventions, e.g. by adequately feeding women during pregnancy in order to prevent chronic diseases in future generations. Additionally, gender-specific mechanisms should be taken into account in future studies.

## Acknowledgments

This work was supported by the European Community [FP7 HEALTH, Project 279281 (BrainAge) to K.F.]. The sponsors had no role in the design and conduct of the study; collection, management, analysis, and interpretation of the data; and preparation, review, or approval of the manuscript.

